# Proteomic analysis of early SASP-associated responses in doxorubicin-treated cardiac fibroblasts

**DOI:** 10.64898/2026.11.10.765715

**Authors:** Angelica M. Riojas, Nazli Gharraee, Zeeshan Hamid, Kathryn W. Dragun, Michael Olivier, Kip D. Zimmerman, Giselle C. Meléndez

**Affiliations:** Department of Cardiology, Wake Forest University School of Medicine, Winston-Salem, NC, USA; Center for Precision Medicine, Wake Forest University School of Medicine, Winston-Salem, NC, USA; Department of Internal Medicine, Section on Molecular Medicine, Wake Forest University School of Medicine, Winston-Salem, NC, USA; Department of Biostatistics and Data Science, Wake Forest University School of Medicine, Winston-Salem, NC, USA

**Keywords:** Anthracyclines, Cardiac fibroblasts, Cellular senescence, Senescence-associated secretory phenotype (SASP), Proteomics, Mitochondrial stress, Anthracycline cardiotoxicity

## Abstract

Anthracycline-induced cardiotoxicity is associated with progressive cardiac dysfunction and fibrotic remodeling, processes in which cardiac fibroblast senescence and the senescence-associated secretory phenotype have been implicated. However, the early secretory responses that precede established senescence following doxorubicin (DOX) exposure remain poorly defined. Primary cardiac fibroblasts isolated from healthy rat hearts were treated with varying concentrations of DOX (0–3 μM) for 24 hours. Media collected at Day 1 were analyzed by untargeted proteomics, while cell viability, proliferation, and senescence-associated β-galactosidase activity were assessed at Days 1 and 7.

Acute DOX exposure reduced cell viability and proliferation at Day 1 without significantly increasing senescence-associated β-galactosidase activity. At Day 7, we observed persistent proliferative arrest and increased β-galactosidase activity with DOX exposure, consistent with delayed senescence. Proteomic analysis of Day 1 conditioned media identified 35 proteins significantly associated with DOX treatment after false discovery rate correction (22 positive and 13 negative associations; P_FDR < 0.05). Functional annotation identified proteins related to extracellular matrix (ECM) regulation, mitochondrial function, cellular stress, cytoskeletal organization, calcium signaling, protease activity, and cell-cycle regulation. Network analysis identified a collagen-centered network (COL1A1, COL1A2, and COL3A1) associated with reduced abundance of ECM proteins and a second network centered on GAPDH, HSPA8, and HSPA9 that included proteins associated with mitochondrial proteostasis, oxidative stress, metabolic adaptation, and structural remodeling.

These findings support that cardiac fibroblasts exposed to DOX mount an early stress-associated response that precedes overt senescence. Because the observed proteomic changes occurred before the changes in SA-β-Gal activity, the Day 1 secretory profile may reflect early fibroblast reprogramming rather than immediate activation of a canonical profibrotic or mature senescence-associated secretory phenotype. This pre-senescent phase may represent a potential window for intervention before persistent senescence and maladaptive cardiac remodeling develop.

## Introduction

Anthracyclines are highly effective chemotherapy agents used to treat hematological malignancies, solid tumors and sarcomas, and triple negative breast cancer^1,2^. However, their clinical utility is limited by dose-dependent cardiotoxicity, which can lead to progressive cardiac function declines^3^. A hallmark of anthracycline-induced left ventricular (LV) dysfunction is interstitial cardiac fibrosis. While many studies have focused on direct anthracycline toxicity on cardiomyocytes, the mechanisms underlying cardiac extracellular matrix (ECM) changes are less defined^4–6^. Cardiac fibroblasts are central regulators of ECM homeostasis, and we previously demonstrated that doxorubicin (DOX, a frequently used anthracycline) directly activates these cells, promoting increased collagen deposition and fibrotic remodeling, findings that are further supported by other groups showing that DOX alters fibroblast function and ECM regulation^4,7–9^.

Previous transcriptomic studies have shown that early DOX responses are dominated by DNA damage and cell-cycle dysregulation rather than activation of canonical fibrotic pathways, despite increased ECM deposition^4,7,8,10^. These findings suggest that mechanisms beyond transcriptional activation of fibrotic pathways may contribute to Dox-induced fibrotic remodeling. Cellular senescence, a state of cell cycle arrest and the development of senescence-associated secretory phenotype (SASP), have emerged as potential mediators of tissue remodeling following DOX exposure^6,11,12^. However, the temporal relationship between early DOX-induced secretome changes, the development of senescence and profibrotic fibroblast remodeling remains poorly defined. Therefore, we investigated the early secretory response of cardiac fibroblasts following acute DOX exposure and evaluated its relationship to subsequent progression to senescence and changes in fibroblast function. Using primary cardiac fibroblasts in vitro, we characterized the early senescence-associated secretory phenotype (SASP) elicited by acute, dose-dependent DOX exposure and evaluated their relationship to subsequent changes in fibroblast phenotype.

## Materials and Methods

### Cardiac fibroblasts isolation

Primary cardiac fibroblasts were isolated from eight-week-old female Sprague Dawley rats (n=4) (Jackson Lab). All animal procedures were performed in accordance with the protocols approved by the Wake Forest School of Medicine Institutional Animal Care and Use Committee (IACUC protocol A22-006). Rats were euthanized by intraperitoneal injection of pentobarbital sodium and phenytoin sodium (100 mg/kg, Euthasol) and subsequent thoracic cavity opening. Hearts were removed and immediately added to a sterile petri dish and placed into a biosafety cabinet where the LV was dissected from the rest of the cardiac structures and immediately placed in ice-cold Krebs–Henseleit buffer (KHB) (2.1 g/L of NaHCO_3_ at pH 7.4). Tissue fragments (∼1 mm² pieces) were washed with KHB and digested with Liberase (Roche) with fibroblasts isolated by primary attachment to cell culture dish. Cell culture were maintained in a humidified incubator under standard conditions (37°C and 5% CO_2_), and medium was replaced every 2 days thereafter.

### Cell culture experiments

All results generated are from primary cardiac fibroblasts at passaged 2 (P2) to ensure consistent cell phenotype. Cells were plated (2 × 10^5^ cells per well for 24 hours to adhere in a Dulbecco’s Modified Eagle Medium (DMEM) supplemented with 1.5% fetal bovine serum (FBS). Media was then replaced DMEM supplemented with 1.5% FBS, F12 and antibiotics (penicillin and streptomycin) containing DOX (0 μM, 0.5 μM, 1 μM, or 3 μM) for an additional 24 hours. For untargeted proteomic analysis, cells were seeded at a density of 2 × 10^5^ cells per well and culture supernatants were collected after 24 h of incubation (Day 1), with three technical replicates for each condition. To assess the long-term effects of DOX on senescence markers in cardiac fibroblasts, additional wells were plated using the same culture conditions. After removal of DOX-containing supernatants, cells were thoroughly washed with phosphate-buffed saline (PBS) and the media was replaced with serum-free DMEM F12 to prevent cell proliferation. Cells were then maintained for an additional 7 days, with a medium changes on days 3 or 4.

### Cell viability assays

Cell viability was assessed using an XTT colorimetric assay. Cells were seeded into 96-well plates (density of 1 × 10^4^ cells; with 5 replicates per group) and all culture conditions and treatment procedures were performed as described above. At experimental timepoint, of XTT reagent (10 uL, 4 mg XTT/4 mL of DMEM; GoldBio), and XTT detection solution (50 μL, 3 mg Phenazine methosulfate in 1 mL of 1X PBS) was added directly to each well. Following a 2-hour incubation at 37 °C, colorimetric absorbance was measured using a Varioskan Lux Plate Reader (Thermo Scientific) at 450 nm. Relative cell viability was determined from absorbance values using a standard curve generated from untreated control wells as previously described^13^.

### Cell proliferation assay

Cell proliferation (plated density of 1 × 10^4^ cells; with 3 replicates per group) was assessed using a 5-ethynyl-2′-deoxyuridine (EdU) incorporation assay by Click-iT Plus EdU Imaging Kit (Life Technologies). The assay was used according to the manufacturer’s instructions. Briefly, EdU was added to the culture medium and incubated (37°C and 5% CO_2_) for 24 hours. Cells were then washed with 1X PBS, fixed with 10% formaldehyde in PBS, and permeabilized with 0.1% Triton X-100. Cells were then subjected to Click-iT reaction using a fluorescent azide labeling solution. Nuclei were stained with 250 μL of DAPI, and images were acquired using under EVOS XL Core Imaging System (Life Technologies). Quantification was performed using ImageJ (Fiji) and cell proliferation was expressed as the proportion of EdU-positive cells (representative of the newly proliferated cells) relative to total number of DAPI-stained nuclei (representative of the total number of cells).

### Senescence-associated β-Galactosidase (SA-β-Gal) Activity

SA-β-Gal staining was assessed using the senescence beta-Galactosidase Staining Kit (#9860, Cell Signalling Technology) according to manufacturer’s instructions (plated density of 1 × 10^4^ cells; with 3 replicates per group). Briefly, cells were washed with PBS (Gibco), fixed with 1X Fixative solution for 15 min at room temperature and then incubated at 4°C overnight. Cells were then stained with the galactosidase staining solution (pH=6) and incubated 37 °C in a dry incubator overnight. Nuclei were counterstained with DAPI, mounted with ProLong™ Gold Antifade Mountant (Thermo Fisher Scientific), and imaged using an EVOS XL Core Imaging System (Invitrogen). Images were analyzed using ImageJ (Fiji), and senescence was quantified as the percentage of SA-β-Gal-positive cells (representative of the number of senescent cells) relative to the total number of DAPI-stained nuclei (representative of the total number of cells per well).

### Untargeted proteomics analysis of secretome

A total of 1μg of each sample was loaded on a PepMap RSLC C18 easy-spray column (3 μM, 100A, 75 μM*15 cm) using Easy-nLC 1200 coupled to an Orbitrap Lumos Mass Spectrometer (Thermo Scientific). The peptides were separated using a 2-hour gradient of Mobile phase A (0.1% Formic acid in 95:5 Water: Acetonitrile) and Mobile Phase B (0.1% Formic acid in 80:20 Acetonitrile: Water). The following gradient was used for peptide elution: 2 to 20% B in 85 minutes, 20% to 95% B in 30 minutes, and final hold of 95% B for next 5 minutes. The mass spectrometer data was acquired in data-dependent acquisition mode, with an MS1 scan range between 375 and 1800 m/z, a resolution of 60,000, and a maximum injection time of 50 ms. MS/MS data acquisition was done in ion trap using CID 35%. All the data acquisition was done using Thermo Scientific Xcalibur software. Raw files were searched with MaxQuant version 1.6.8.0. as label free quantification. Match between runs was set to false, minimum peptide length was set to 7, decoy mode was set to revert, and contaminants were included in the search. The digestion rule was set to trypsin/p. Carbamidomethylation was enabled as a fixed modification. Variable modifications are N-terminal acetylation and oxidation of methionine. Protein and peptide identification FDR was set to 1%.

### Statistical Analysis

#### Cardiac Fibroblast Phenotype Measures

Global differences in cell viability, proliferation, and the percentage of β-gal positive cells across DOX treatment groups (0, 1, 3 µM and time points (24 hours and 7 days) were assessed by 2-Way ANOVA. For each outcome, contribution of DOX dose, time and their interaction (DOX dose x time) to overall variance was determined. Pre-specified Sidak’s multiple comparisons with corrections were computed for all doses compared between Day 1 and Day 7. Data are presented as individual biological replicate values with mean ± standard deviations and P < 0.05 were considered statistically significant.

#### Statistical Analysis of Proteomics Data

The raw protein intensities were normalized using total protein normalization by the median-of-ratios method^14^. The normalized protein intensities were then log-transformed. Proteins were sorted into two groups: those with missing values were retained for logistic regression, while those with complete data were retained for linear regression. Proteins with complete data were used to test for differences in magnitude (abundance) using linear regression. Proteins with missing values present were used to test for differences in detection (presence/absence) by logistic regression. For the logistic regression, missing values were set to 0 and all other values were set to 1. Association testing with the logistic regression was computed to test for increased presence or absence of a protein with dose. The linear regression tested for association between the normalized protein abundance and dose. To ensure reproducibility of an initial round of proteomics, the proteomics experiment was replicated from start to finish. To specifically identify proteins that were consistently significant and in the same direction across both of the repeated experiments an inverse variance based meta-analysis method was used^15^. To adjust for multiple comparisons, an Benjami-Hochberg False Discovery Rate (P_FDR_) correction was applied^16^.

Those proteins meeting an P_FDR_ < 0.05 were retained for network analysis. Proteins were submitted to STRING to identify protein-protein interactions as a way of identifying functionally related sets of proteins that were altered between groups. “Text mining” and “Neighborhood” interactions were disabled to focus on experimentally determined interactions. STRING networks were generated with all interaction sources and medium confidence (0.4) threshold with a k-means clustering threshold of 3.

Proteins significantly associated with DOX concentration were annotated using Gene Ontology terms obtained through Ensembl BioMart (ensemble.org). For each protein, the presence or absence of annotations related to senescence, cell-cycle regulation, collagen organization, cell structure, fibrosis, protease activity, mitochondrial function, and calcium signaling was recorded. Because individual proteins could be associated with multiple terms, these annotations were not mutually exclusive and were visualized as a functional annotation heatmap in Figure 2. Proteins were also assigned to a primary cardiovascular functional category based on literature review, with detailed justification and supporting references provided in Table 1 and Supplemental Table S1.

**Table 1.**
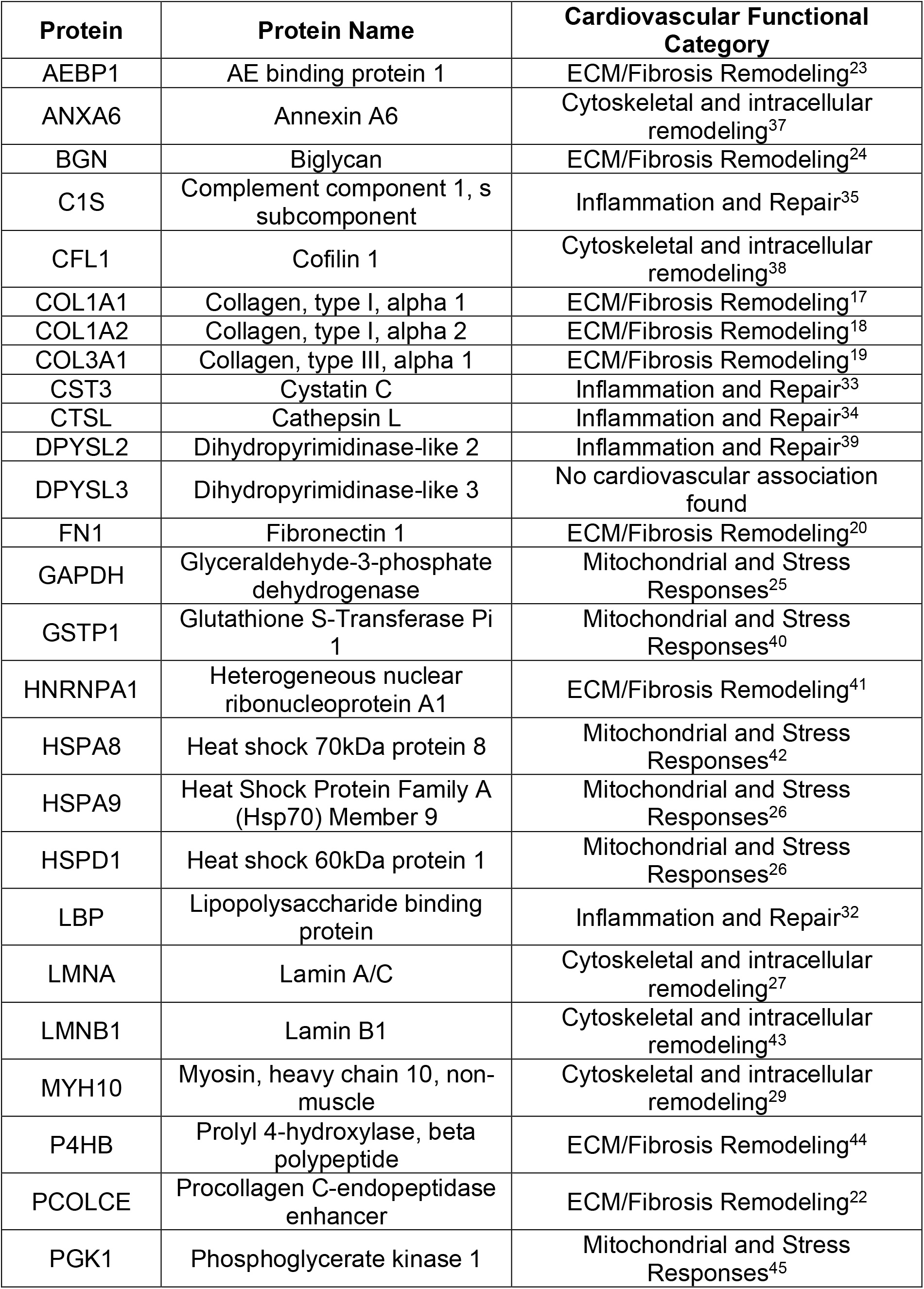
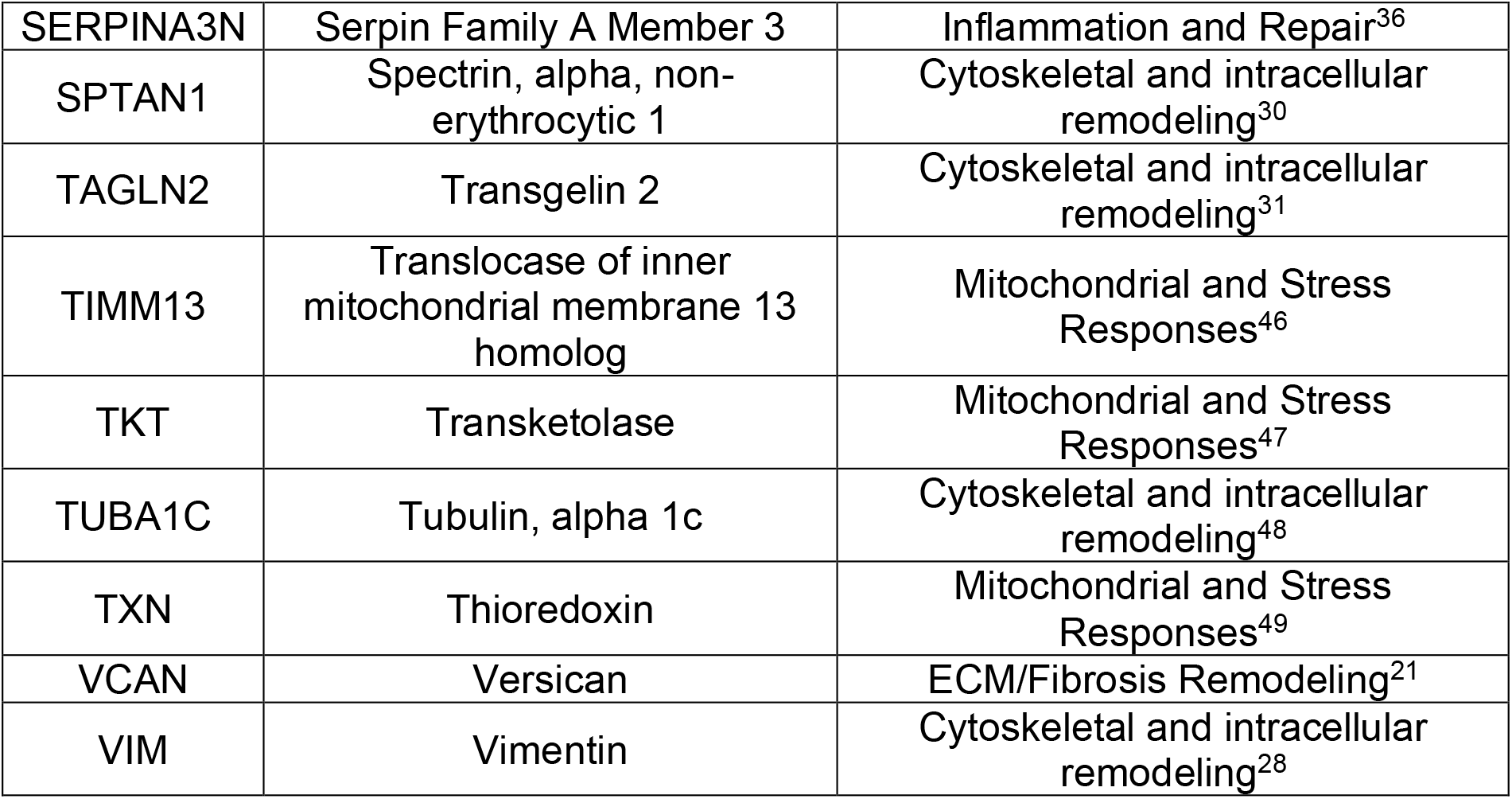
Cardiovascular functional classification of proteins significantly associated with DOX concentration. Proteins identified by proteomics of media from DOX treated cells were assigned to one primary cardiovascular functional category based on their predominant role reported in the literature. Because proteins may participate in multiple biological processes, these categories were used to facilitate interpretation and are not mutually exclusive. Detailed evidence type, study findings, cardiovascular relevance, and PubMed identifiers supporting each classification are provided in Supplemental Table S1. DOX: Doxorubicin.

## Results

### Cardiac fibroblasts undergo early growth arrest followed by delayed senescence marker expression after DOX exposure

To determine whether DOX exposure altered the number of viable cardiac fibroblasts over time, estimated viable cell numbers were analyzed across doses and days. Viable cell number was significantly influenced by DOX dose and time, with a significant interaction between these factors (day: F_(1,24)_ = 105.26, p < 0.0001; dose: F_(2,24)_ = 149.24, p < 0.0001; day-by-dose interaction: F_(2,24)_ = 16.62, p < 0.0001). At Day 1, viable cell number did not differ between untreated and 1 μM DOX-treated cells (0 μM: 9.47 ± 1.37 × 10^3^ vs. 1 μM: 10.45 ± 0.50 × 10^3^ cells; p = 0.84). In contrast, viable cell number was reduced following 3 μM DOX (4.54 ± 0.36 × 10^3^ cells) compared with both untreated and 1 μM DOX-treated cells (both p < 0.0001; Figure 1A).

**Figure 1.**
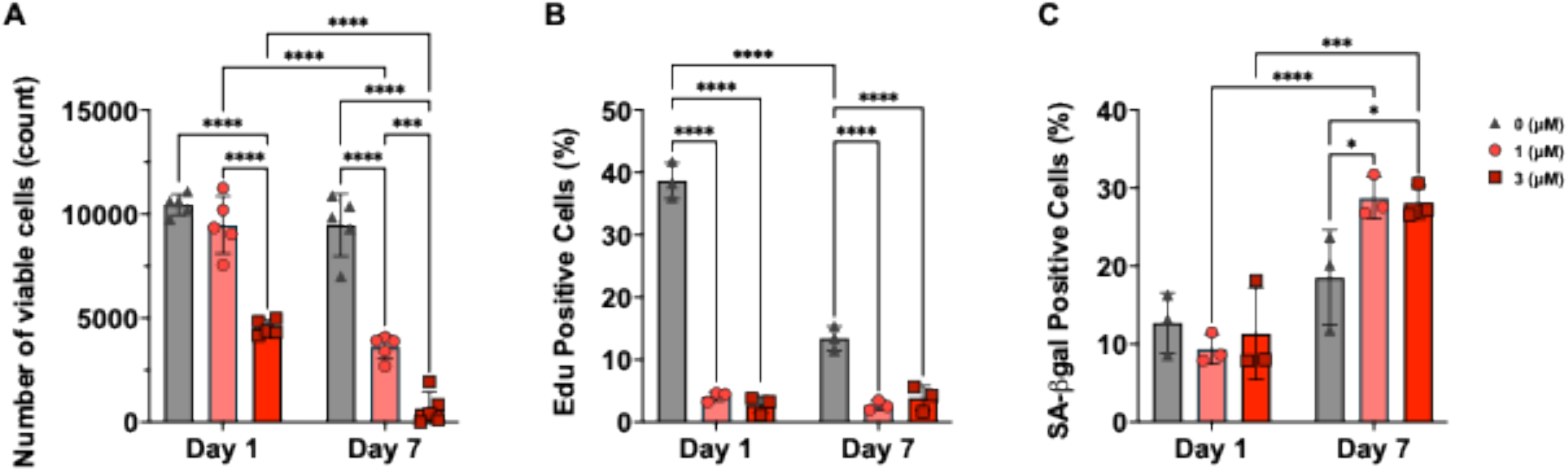
Day 1 and Day 7 cardiac fibroblast characterization with dox treatment; early SASP. Cell culture measures of A) cell number from XTT, B) EdU positive cells, and C) beta-gal positive cells are shown for controls (0 uM, grey triangles) and increasing concentrations of dox (1 uM, pink circles; 3 uM red squares). Statistical analysis was performed using 2 way ANOVA with Sidak’s multiple comparison test, with n=3 technical replicates in each graph and significant p values shown above brackets as *for each comparison with significance defined as p <0.05.

At Day 7, viable cell number was reduced in both DOX-treated groups compared with untreated controls (0 μM: 9.48 ± 1.50 × 10^3^ cells vs. 1 μM: 3.60 ± 0.57 × 10^3^ cells, p < 0.0001; vs. 3 μM: 0.66 ± 0.79 × 10^3^ cells, p < 0.0001). Viable cell number was also lower following 3 μM compared with 1 μM DOX (p = 0.0009). From Day 1 to Day 7, viable cell number declined within the 1 μM and 3 μM groups but remained unchanged in untreated controls (0 μM: p = 0.85, both 1 μM and 3 μM: p < 0.0001). Together, these findings demonstrate a dose- and time-dependent reduction in viable cell number following DOX exposure.

We next assessed EdU incorporation to determine whether reduced viable cell number was accompanied by impaired proliferation. DOX exposure induced a rapid and persistent reduction in cardiac fibroblast proliferation that varied with dose and time (day: F_(1,12)_ = 101.15, p < 0.0001; dose: F_(2,12)_ = 318.11, p < 0.0001; day-by-dose interaction: F_(2,12)_ = 97.68, p < 0.0001). At Day 1, EdU incorporation was reduced following both 1 μM and 3 μM DOX compared with untreated controls (0 μM: 38.64 ± 2.89% vs 1 μM: 4.14 ± 0.77%; p < 0.0001 and 3 μM DOX: 2.77 ± 1.40%; p < 0.0001, Figure 1B). A similar pattern persisted at Day 7, with lower EdU incorporation following 1 μM (2.68 ± 0.75%; p = 0.0001) and 3 μM DOX (3.87 ± 2.02%; p = 0.0004) compared with untreated controls (13.43 ± 1.96%).

A similar pattern persisted at Day 7, with lower EdU incorporation following 1 μM and 3 μM DOX compared with untreated controls (0 μM: 13.43 ± 1.96% vs 1 μM (2.68 ± 0.75%; p = 0.0001 and 3 μM: 3.87 ± 2.02%; p = 0.0004). EdU incorporation did not differ between the 1 and 3 μM groups at Day 1 or Day 7 (1 μM p = 1.00, 3 μM p = 1.00). From Day 1 to Day 7, EdU incorporation declined in untreated cells but did not change significantly within either DOX-treated group (0 μM: p < 0.0001, 1 μM p = 1.00, 3 μM p = 1.00). Thus, DOX-induced proliferative arrest was established by Day 1 and persisted through Day 7, without additional suppression at 3 μM compared with 1 μM DOX.

Finally, we measured SA-β-gal activity at Days 1 and 7 to determine when a senescence-associated phenotype emerged following acute DOX exposure and whether it was evident at the early time point in a dose-dependent manner. SA-β-Gal activity increased over time following DOX exposure, with the temporal response differing by dose whereas the main effect of DOX dose was not significant (day: F_(1,12)_ = 51.65, p < 0.0001; day-by-dose interaction: F_(2,12)_ = 4.54, p = 0.034; dose: F_(2,12)_ = 1.70, p = 0.22). At Day 1, SA-β-Gal positivity was similar among untreated and 3 μM DOX-treated cells (0 μM: 12.69 ± 3.88% vs 1 μM: 9.34 ± 1.92% and 3 μM: 11.32 ± 5.88%; all p ≥ 0.999; Figure 1C). At Day 7, SA-β-Gal positivity was increased in 1 μM DOX and 3 μM DOX compared to untreated cells (0 μM: 18.52 ± 6.12% vs 1 μM: 28.70 ± 2.66%, p = 0.0323 and 3 μM: 28.11 ± 2.15%, p = 0.0445); comparisons between 1 μM DOX and 3 μM DOX doses were not significant (p ≥ 0.999). However, SA-β-Gal positivity increased from Day 1 to Day 7 within the 1 μM and 3 μM groups (p = 0.0048), but not within untreated controls (0 μM: p = 0.83; 1 μM: p = 0.0014; 3 μM: p = 0.0048). Together, these findings indicate that DOX-induced proliferative arrest preceded the emergence of a detectable senescence-associated phenotype.

### Acute DOX exposure suppresses extracellular matrix proteins while increasing mitochondrial stress and intracellular remodeling proteins

To evaluate the early secretory response to acute DOX exposure, we performed untargeted proteomic analysis of conditioned media collected 24 hours after treatment. A total of 35 proteins were significantly associated (P_FDR_ < 0.05) with increasing DOX concentration (22 positively associated and 13 negatively associated; Figure 2). Functional annotation using Gene Ontology information identified proteins associated with biological processes related to senescence, cell cycle regulation, collagen organization, cell structure, fibrosis, protease activity, mitochondrial function, and calcium signaling (Figure 2). Proteins were additionally assigned to one of four cardiovascular functional categories based on literature review: (1) ECM/fibrosis remodeling, (2) mitochondrial and stress responses, (3) cytoskeletal and intracellular remodeling, and (4) inflammation and repair (Table 1; Supplemental Table S1).

**Figure 2.**
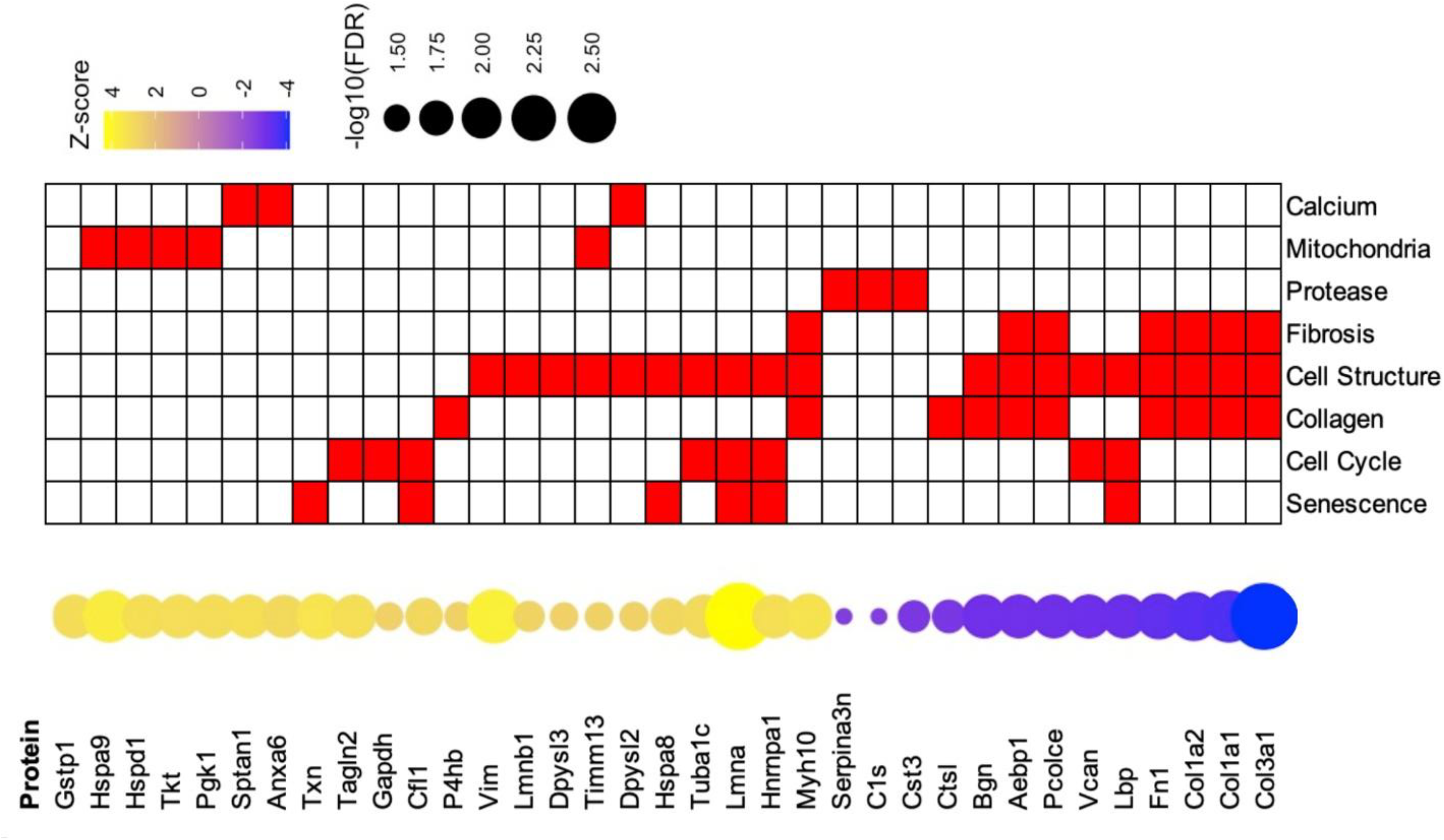
Heatmap of early Day 1 SASP profile and functional summary. Proteins identified in Day 1 conditioned media are listed next to circle with Z-score indicating upregulation (yellow) or down regulation (blue) in relation to increasing dox dose (0,1,3 uM) with circle size corresponding to significance by the –log10(FDR). Squares in heatmap indicate functional association by classification of presence (red) or absence (white).

Among the proteins demonstrating the greatest negative associations with increasing DOX concentration were several ECM-associated proteins, including COL3A1, COL1A1, COL1A2, FN1, VCAN, PCOLCE, AEBP1, and BGN (Figure 2). In contrast, proteins demonstrating the greatest positive associations with DOX concentration included HSPA9, TXN, HSPD1, TKT, PGK1, GSTP1, HSPA8, TIMM13, and GAPDH. Additional proteins showing positive associations were structural and cytoskeletal regulators such as LMNA, VIM, MYH10, SPTAN1, TAGLN2, and TUBA1C. Proteins associated with inflammation and repair, including LBP, CTSL, CST3, C1S, and SERPINA3N, were predominantly negatively associated with increasing DOX concentration (Figure 2; Table 1).

To place the proteomic findings into a cardiovascular context, proteins were organized according to previously reported roles in cardiac remodeling, fibrosis, mitochondrial adaptation, cellular stress responses, inflammation, and tissue repair (Table 1; Supplemental Table S1). STRING network analysis identified physical interactions among 31 of the 35 significant proteins and organized them into two major functional clusters (Figure 3). The first cluster was centered on COL1A1, COL1A2, and COL3A1 and consisted primarily of proteins involved in ECM organization, collagen processing, and fibrosis-related pathways. The second cluster was centered on GAPDH together with HSPA8 and HSPA9 and contained proteins associated with mitochondrial proteostasis, oxidative stress responses, metabolic regulation, and cellular stress adaptation. Several proteins, including VCAN, LBP, C1S, SERPINA3N, and TIMM13, demonstrated significant dose-associated changes but were not incorporated into either major interaction cluster. Together, these findings indicate that increasing DOX exposure is associated with coordinated reductions in ECM- and repair-associated proteins alongside increased abundance of proteins involved in mitochondrial stress responses and cellular structural remodeling.

**Figure 3.**
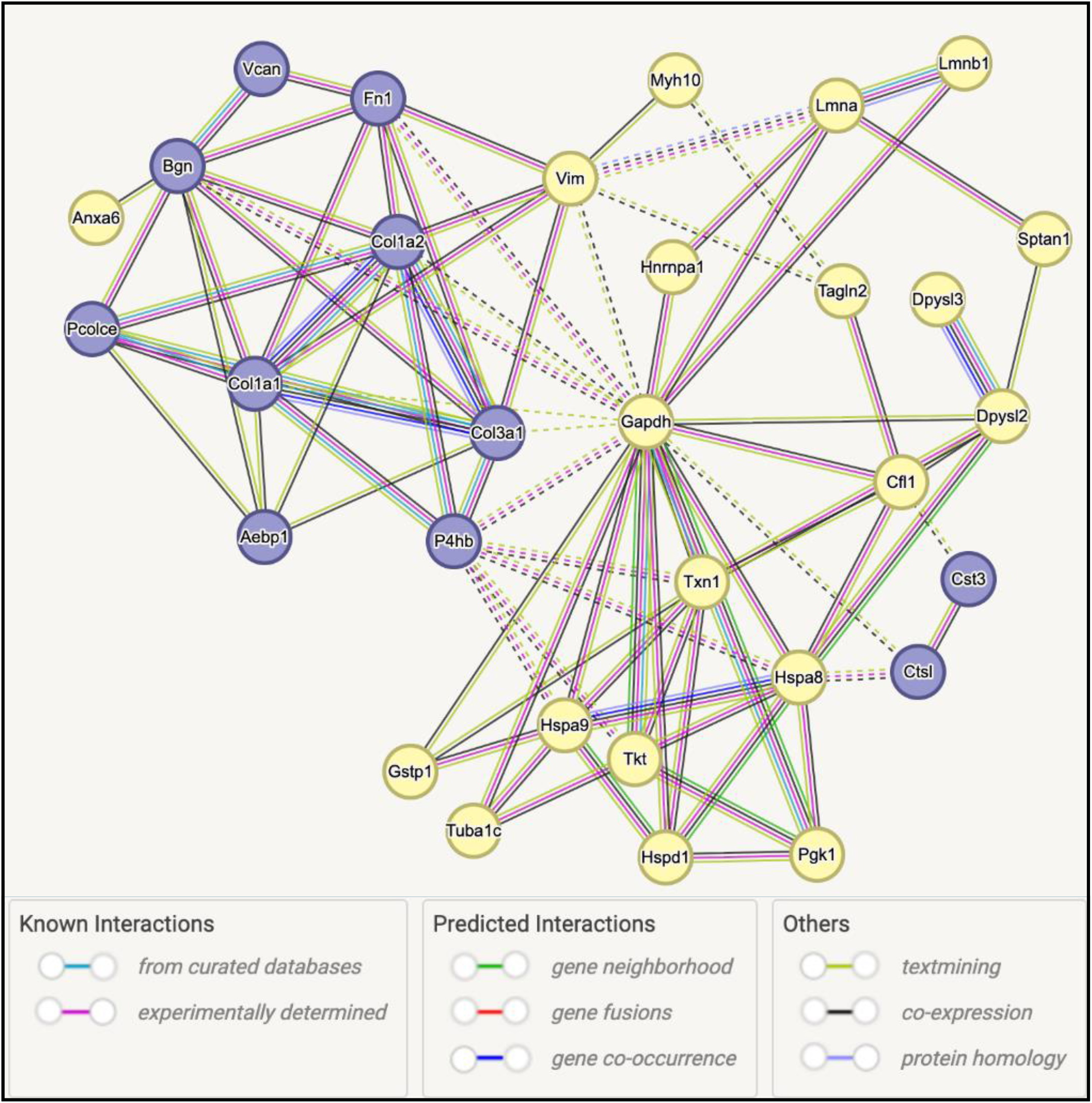
String analysis; protein interactions with clustering. Significant proteins identified in Day 1 conditioned media in relation to increasing dox dose (0,1,3 uM) are shown from STRING (https://string-db.org/) protein associations with clusters colored by grouping (red, green, blue). Proteins are outlined by a circle corresponding to Z-score (yellow-positive, blue-negative). Line colors connecting proteins are colored by type and source of interaction annotation, with dotted line indicating links between clusters determined by k-means.

## Discussion

Previous studies have established that DOX directly affects cardiac fibroblasts and contributes to ECM remodeling independent of cardiomyocyte injury^3,4,8,9^. In the present study, acute DOX exposure reduced fibroblast viability and proliferation within 24 hours but did not induce a robust senescent phenotype at Day 1. In contrast, persistent proliferative arrest and increased senescence-associated β-galactosidase activity were evident after seven days of recovery, suggesting that growth arrest and cellular stress precede the development of overt senescence (Figure 1). Together, these findings indicate that early fibroblast response to DOX involves cellular stress and proliferative arrest that precedes the development of overt senescence.

We identified a cluster of ECM proteins around collagen proteins COL1A1, COL1A2, and COL3A1. Many of the proteins within this network, including FN1, VCAN, PCOLCE, AEBP1, and BGN, have established roles in fibroblast activation, ECM remodeling, fibrosis, and adverse cardiac remodeling (Table 1)^17–24^. Rather than supporting immediate activation of a classical profibrotic program, the decreased abundance of these proteins suggests that acute DOX exposure initially suppresses or reorganizes ECM-regulatory pathways. These findings are consistent with prior observations that early DOX responses are dominated by cellular stress and injury pathways rather than established fibrotic remodeling or senescence^10^.

In contrast, proteins associated with mitochondrial homeostasis, oxidative stress responses, and metabolic adaptation were predominantly increased following DOX exposure. The second major cluster in the network was centered on GAPDH together with the molecular chaperones heat shock protein 8 (HSPA8) and HSPA9 and included multiple proteins involved in mitochondrial proteostasis, redox regulation, and stress adaptation. Previous studies in murine models of cardiac aging and ischemia have implicated the proteins in this cluster with mitochondrial quality control, cardioprotection, oxidative stress defense, and adaptation to cellular injury^25,26^. Given the established role of DOX in disrupting mitochondrial function through oxidative stress and altered calcium handling, the coordinated activation of this network supports mitochondrial stress as an early feature of fibroblast injury that precedes overt senescence.

Importantly, the mitochondrial stress-response network also contained many of the proteins assigned to the cytoskeletal and intracellular remodeling cluster, including LMNA, VIM, MYH10, SPTAN1, TAGLN2, CFL1, and TUBA1C. Several of these proteins showed some of the strongest positive dose-associated changes observed in the dataset. As summarized in Table 1, these proteins have been shown in human and murine cardiac studies to regulate cellular architecture, mechano-transduction, migration, contractility, and structural adaptation in cardiovascular tissues^27–31^. The localization of these proteins within the same interaction network as mitochondrial stress-response proteins suggests that structural remodeling and stress adaptation are likely tightly coupled components of the early DOX response. Rather than immediately increasing ECM production, surviving fibroblasts appear to undergo intracellular structural remodeling that may influence cellular mechanics, communication, and subsequent remodeling responses.

Proteins previously associated with inflammation and tissue repair in human cell culture and rodent models were also predominantly reduced following DOX exposure. Reduced abundance of LBP, CST3, CTSL, C1S, and SERPINA3N suggests reorganization of pathways involved in innate immune signaling, ECM turnover, complement activation, and post-injury repair^32–36^. Notably, this pattern did not support broad activation of inflammatory pathways. Instead, the findings support a model in which extracellular repair-associated signaling is altered during the acute injury response. These observations further support interpretation of the Day 1 secretory profile as a stress-associated response rather than a mature senescence-associated secretory phenotype. Collectively, these findings suggest that acute DOX exposure initially suppresses ECM and repair-associated pathways while activating mitochondrial stress responses and intracellular structural remodeling within surviving fibroblasts. Because these changes occur before robust β-gal induction, the Day 1 secretory profile is more appropriately interpreted as an early stress-associated or pre-senescent response rather than an established SASP. From a clinical perspective, these findings raise the possibility that cardiac fibroblast dysfunction may occur before a profibrotic phenotype and widespread senescence become apparent, potentially identifying a window for intervention targeting mitochondrial stress or maladaptive fibroblast reprogramming.

Several limitations should be acknowledged. First, the study utilized only female rat cardiac fibroblasts and may not fully reflect responses that occur in males. Second, the *in vitro* system does not account for interactions among fibroblasts, cardiomyocytes, endothelial cells, and immune cells that contribute to cardiac remodeling and potentially senescence induction. Third, proteomics of cell culture media cannot definitively distinguish actively secreted proteins from intracellular proteins released during cell injury or death. Although the coordinated network structure argues against purely nonspecific protein leakage, future studies incorporating cellular proteomics and targeted validation will be necessary to address this limitation. Finally, the study examined a single acute exposure treatment regimen, whereas clinical DOX treatment involves repeated dosing and complex multicellular responses. Future studies incorporating co-culture systems, *in vivo* validation, single-cell approaches, and direct assessment of mitochondrial function will be necessary to determine how these early stress-response pathways contribute to delayed senescence and DOX-associated cardiac remodeling.

## Declaration of Generative AI and AI-assisted technologies in the writing process

During the preparation of this work the author(s) used Microsoft Copilot to outline and improve paragraph structure. After using this tool/service, the author(s) reviewed and edited the content as needed and take(s) full responsibility for the final content of the publication.

## Author contributions

G.C.M. conceptualized and designed the study, performed experimental work, contributed to data analysis, and secured funding. M.O. designed and oversaw all aspects of the proteomics experiments and analyses. N.G. contributed to the experimental design, conducted experimental studies, and performed cell culture data analysis. Z.H. conducted proteomic experimental studies, A.M.R. conducted proteomic data analyses, visualization, and validation, and prepared the original manuscript draft. K.W.D. contributed to original manuscript draft. K.D.Z conducted proteomic data analyses, visualization, and validation. All authors reviewed and approved the final manuscript.

## Data Availability

The mass spectrometry proteomics data have been deposited in the ProteomeXchange Consortium via the PRIDE partner repository at EMBL-EBI under accession number PXD083374. Reviewer access details are available upon request or provided with the submission materials.

